# Identification of the Calmodulin-dependent NAD^+^ kinase sustaining the elicitor-induced oxidative burst in plants

**DOI:** 10.1101/521658

**Authors:** Elisa Dell’ Aglio, Cécile Giustini, Alexandra Kraut, Yohann Couté, Christian Mazars, Michel Matringe, Giovanni Finazzi, Gilles Curien

**Affiliations:** Univ. Grenoble Alpes, CNRS, CEA, INRA, BIG-LPCV, 38000 Grenoble, France; Department of Botany and Plant Biology, University of Geneva, 1211 Geneva, Switzerland; Univ. Grenoble Alpes, CEA, INSERM, BIG-EdyP, 38000 Grenoble, France; Laboratoire de Recherche en Sciences Végétales, Université de Toulouse, CNRS, UPS, 24, Chemin de Borde-Rouge, Auzeville, BP 42617, 31326 Castanet-Tolosan, France

**Keywords:** NAD kinase, Calmodulin, Calcium, NADP^+^, zeta toxin, flagellin, *Arabidopsis thaliana*

## Abstract

NADP(H) is an essential cofactor ofmultiple metabolic processes in all living organisms. While NADP+ production in plants has long been known to involve a Calmodulin (CaM)/Ca^2+^-dependent NAD^+^ kinase, the nature of the enzyme catalyzing this activity has remained enigmatic, as well as its role in plant physiology. Here, we identify an Arabidopsis P-loop ATPase (Atlg04280) with a bacterial type II zeta toxin domain, that catalyzes NADP^+^ production upon binding of CaM/Ca^2+^ to a domain located in its N-terminal region. The encoded protein (NADKc-1) is associated with the mitochondria and amplifies the elicitor-induced oxidative burst in Arabidopsis leaves representing the missing link between calcium signalling and metabolism in the response to pathogen elicitor. By analysis of various plants and algae, we show that NADKc is well conserved in the plant lineage and present in basal plants. Our data allows proposing that the CaM-dependent NAD kinase activity is only found in photosynthetic species carrying NADKc-1 related proteins, which would represent the only proteins harboring CaM-dependent NAD kinase activity in plants and algae.

## Introduction

As sessile organisms, plants have evolved fast means to detect the presence of pathogens and to react by orchestrating defense barriers. Early reaction to pathogen aggression requires recognition of microbial-derived elicitors commonly known as MAMPs (Microbe-Associated Molecular Patterns, for reviews see (Choi and Klessig, 2016; Newman et al., 2013)), including oligogalacturonides, chitin, flagellin22 (flg22) and proteins coded by AVR genes (Garcia-Brugger et al., 2006). MAMPS are recognized by pattern recognition receptors (PRRs), *i.e.* receptor-like kinases (RLKs) or receptor-like proteins (RLPs) localized at the plasma membrane (Newman et al., 2013). MAMPs recognition triggers early responses such as extracellular Ca^2+^ influxes (Tavernier et al., 1995), Ca^2+^ release from internal pools through IP3- and cADPR-regulated Ca^2+^ channels (Lamotte et al., 2004; Lecourieux et al., 2002) and the activation of a signaling cascade involving MAP kinases (Jonak et al., 2002; Nakagami et al., 2005).

A very early event after pathogen recognition (3-10 minutes) is the induction of an apoplastic oxidative burst (ROS burst) generated by plasma membrane NADPH oxidases known as RBOHs (for Respiratory Burst Oxidase Homologs) (Torres and Dangl, 2005). ROS production mediates stomatal closure (Munemasa et al., 2015) and might be involved in other long-distant signaling processes. RBOH oxidase activity is dependent on Ca^2+^ binding to their EF-hand domains and is stimulated by phosphorylation by CPKs (Ca^2+^-dependent protein kinases), *i.e.* Ser/Thr protein kinases that are first activated by Ca^2+^ (Adachi and Yoshioka, 2015; Kaur et al., 2014).

In parallel to these well-studied processes, other metabolic changes have been reported during pathogen response. For instance an increase of NADP(H) is observed in response to MAMP treatment (Harding et al., 1997; Pugin et al., 1997). The consequential increase in NADPH cytosolic concentration is supposed to fuel RBOH proteins and sustain the ROS burst (Grant et al., 2000; Torres and Dangl, 2005; Won-Gyu et al., 2017). Because the NADP^+^ level is low in the cytosol (Harding et al., 1997) it has been suggested that plant Calmodulin (CaM)/Ca^2+^-dependent NAD kinase (Anderson and Cormier, 1978; Li et al., 2018) might be a downstream target of Ca^2+^ fluxes, enhancing ROS production by increasing the cytosolic NADP^+^ level. The produced NADP^+^ may be converted to NADPH (the substrate of NADPH oxidase) by the isocitrate dehydrogenase (Mhamdi et al., 2010) and by the reducing branch of the oxidative pentose phosphate pathway, which is stimulated during the pathogen response (Pugin et al., 1997; Rojas et al., 2014; Scharte et al., 2009).

Despite the demonstration that most (~70-90%) of NAD^+^ kinase activity in plant extracts is Calmodulin (CaM)/Ca^2+^-dependent (Delumeau et al., 2000a; Dieter and Manne, 1984; Simon et al., 1982) and present in a wide range of plant species (Anderson et al., 1980; Anderson and Cormier, 1978; Muto and Miyachi, 1977), the gene coding this protein was never identified (Delumeau et al., 1998; Delumeau et al., 2000b; Waller et al., 2010). The plastidial NADK2 was shown to bind CaM *in vitro* in a Ca^2+^-dependent way (Dell’Aglio et al., 2013; Turner et al., 2004) but its activity did not require CaM binding (Turner et al., 2004). This lack of knowledge of the identity of the plant CaM-dependent NAD kinase, has prevented so far a thorough characterization of its role in the production of the ROS burst and in plant pathogen responses.

Here, we characterize a new *A. thaliana* CaM/Ca^2+^-dependent NAD^+^ kinase that displays all the properties of the elusive enzyme. We show that this NAD^+^ kinase, which is associated with mitochondria, is involved in the plant response to the bacterial elicitor flagellin22.

## Results and discussion

### NADKc-1 is a new CaM/Ca^2+^-dependent NAD^+^ kinase

As a first step toward isolation of the high CaM/Ca^2+^-dependent NAD^+^ kinase activity in *A. thaliana*, we enriched a seedling protein fraction containing this activity by applying four purification steps (see Methods and Fig. S1). In the last step of the procedure, proteins binding the CaM matrix in the presence of Ca^2+^ were released by adding an excess of the Ca^2+^ chelator EGTA. We then used mass spectrometry-based proteomics to identify proteins that were strongly enriched in the EGTA elution compared to the Ca^2+^-containing washing steps (Supplemental Table S1). We reasoned that putative candidates should display the following characteristics: *i)* have a molecular weight between 50 and 60 kDa (Delumeau et al., 2000b); *ii)* be annotated as ATP-binding proteins; *iii)* have a predicted CaM-binding site. Our analysis revealed only one protein - coded by the Atig04280 gene - that fulfilled all these criteria.

To confirm its CaM/Ca^2+^-dependent NAD^+^ kinase activity, we expressed the full-length recombinant protein coded by Atig04280 in *E. coli* with an N-terminal His-tag. Activity measurements with a partially purified enzyme revealed absence of NAD kinase activity in the absence of *A. thaliana* (At) CaM1/Ca^2+^. However, NAD kinase activity was revealed within seconds upon addition of both AtCaM1 and Ca^2+^ (Fig. 1A). NAD kinase activity was suppressed by EGTA and restored by the addition of an excess of Ca^2^+, thus enzyme activation is an all-or-none, reversible process (Fig. 1A). Based on these results, we named the Atig04280 gene product NADKc-1, for NAD^+^ kinase CaM-1.

**Figure 1.**
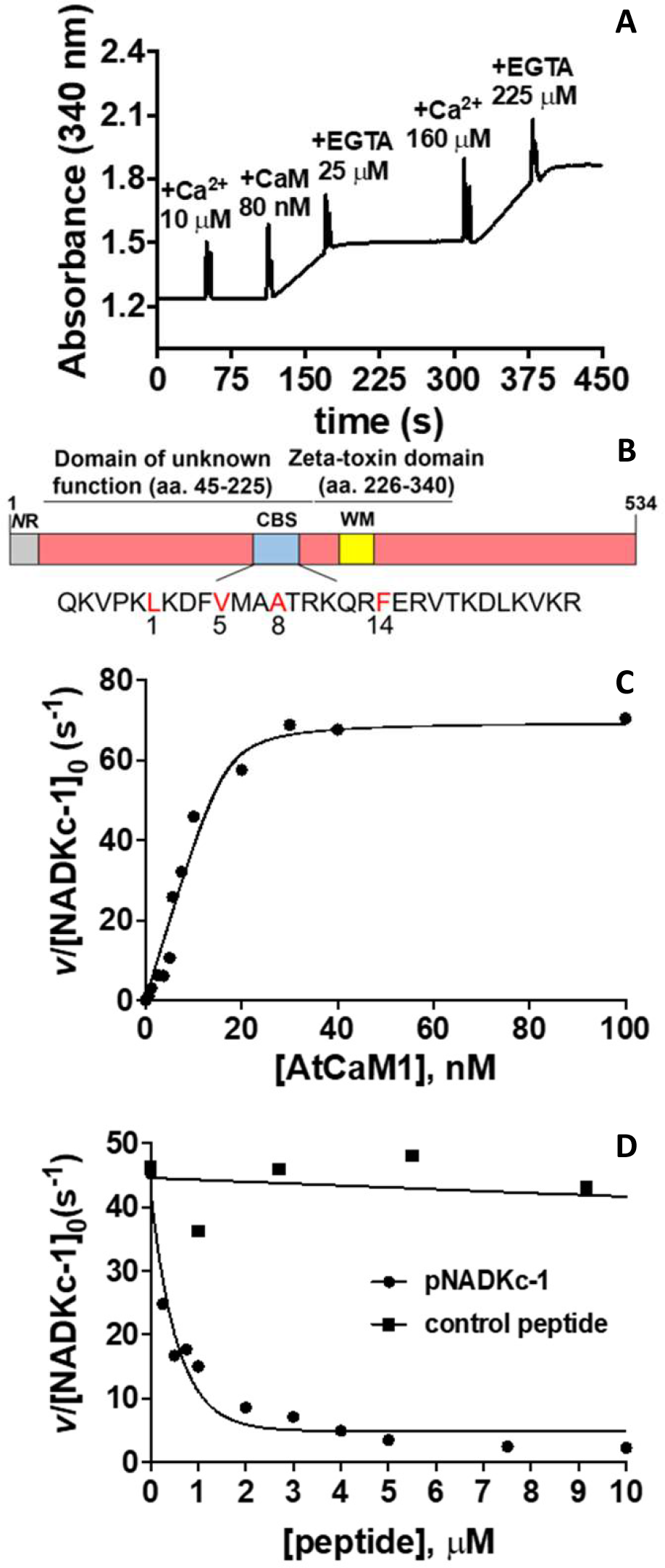
A CaM-dependent NAD kinase identified in Arabidopsis. **(A)** NAD^+^ kinase activity in an *E. coli* bacterial extract expressing theNADKc-1 protein(gene Atlg04280). Ca^2+^, AtCaM1 and EGTA were added at different times, as indicated in the graph. **(B)** Schematic representation of theNADKc-1 primary sequence.NR:N-terminal Region with internal hydrophobic region; CBS: putative conserved CaM-binding site; WM: Walker A motif (ATP binding site). **(C)** Affinity of NADKc-1 recombinant protein for CaM: Activity of the purifiedNADKc-1 recombinant protein after denaturation in urea and subsequent refolding was measured in the presence of 50 μM Ca^2+^ and as a function of [AtCaM1]. Experiments were performed in triplicate and data shown are from one representative experiment. Binding data were analyzed assuming tight-binding. K_d_ value for AtCaM1 binding varied from 0.6 to 1 nM. **(D)** Inhibition of NADKc-1 activity by competition with the putative CaM-binding site (black dots). Black squares correspond to results obtained with a negative control peptide, which does not bind CaM.

The primary sequence of NADKc-1 presented in Fig. 1B contains: *i)* an N-terminal Region (NR) which is predicted to contain a transmembrane helix (amino acids: 24-45, Fig S2); *ii)* a domain of unknown function with no predicted annotation (amino acids 46-225) - this domain, however, contains a conserved CaM-binding site (CBS, Fig. S3A); and *iii)* a C-terminal kinase domain (amino acids 226-340) annotated as a “zeta toxin domain” which is predicted to contain a conserved P-loop for ATP binding (Walker A motif, WM, amino acids: 236-250, Fig. S3B).

To optimize expression levels and improve solubility of the protein in *E. coli*, we removed the first 38 amino acids at the N-terminus - containing a stretch of hydrophobic amino acids. The shorter version, 6HIS-Δ38NADKc-1 was purified by Ni-NTA affinity chromatography (Fig. S4, lane 3). Activity assays with saturating AtCaM1/Ca^2+^ concentrations revealed the NAD^+^ kinase activity of NADKc-1 to be specific toward NAD^+^, as no activity could be detected with NADH or deamido-NAD^+^ (NAAD) (Supplemental Table S2). Like most P-loop containing kinases (Das et al., 2013), the enzyme displayed broad specificity for the phosphoryl donor, as ATP, CTP, GTP and UTP could be used indifferently and produced similar efficiencies (Supplemental Table S2). The enzyme catalytic constant with CTP or ATP was close to 40 s^−1^. Thus, the NADKc-1 catalytic constant is about 10-fold higher than that reported for plant CaM/Ca^2+^-independent NAD^+^ kinases and other NAD^+^ kinases from bacteria and animals (0.5-7 s^−1^, (Apps, 1968; Berrin et al., 2005; Chai et al., 2006; Lerner et al., 2001; Love et al., 2015; Turner et al., 2004; Zerez et al., 1987)).

### Characterization of the NADKc-1-CaM interaction

To characterize the interaction of NADKc-1 with CaM, the recombinant enzyme was further purified. After denaturation of the enzyme in urea and purification on Ni-NTA sepharose in the presence of urea, the protein was refolded by rapid urea dilution (see Methods). As shown in Fig S4, lanes 4-5, the refolded protein produced a single band on an SDS-PAGE gel. The refolded NADKc-1 protein had an increased catalytic constant (70 s^−1^) compared to the partially purified enzyme (40 s^−1^, Supplemental Table S2) and high affinity for CaM (*k*_d_ = 0.6-1 nM, Fig. 1C), similar to the value of 0.4 nM reported elsewhere (Delumeau et al., 2000b).

The CaM target database search tool (Crivici and Ikura, 1995; Yap et al., 2000) predicts a CBS in the N-terminal NADKc-1 domain (amino acids: 168-200, Fig. 1B). This site contains a set of four hydrophobic amino acids (Ll72, Vl76, Al79, F185) forming a “type 1-5-8-14” CBS (Crivici and Ikura, 1995; Yap et al., 2000). In support of a specific role for this domain, it was found to be conserved across homologous proteins in other plant species (Fig. S3A).

To verify that the NADKc-1 N-terminal domain is involved in CaM-binding, we measured NADKc-1 activity in the presence of a synthetic peptide containing the putative NADKc-1 CaM-binding sequence (amino acids 167-196, Fig. 1B) in a competitive assay. As shown in Fig. 1D, the presence of the putative NADKc-1 CaM-binding peptide decreased the stimulation of NADKc-1 by AtCaM1, as expected if AtCaM1, trapped by the peptide in excess, was no longer available for NADKc-1 activation. The reduction in reaction rate was hyperbolically related to the peptide concentration (IC50 =0.5 μM, Fig. 1D). To ensure that inhibition was not simply due to the presence of an excess of any peptide in the reaction, another unrelated peptide from the Tic32 protein, which cannot specifically bind AtCaM1 (Dell’Aglio, 2013), was also tested. An excess of this control peptide had no effect on NAD^+^ kinase activity (Fig. 1D).

These data suggest that the NADKc-1 peptide identified plays a major role in the CaM/Ca^2+^-dependent activation of the NADKc-1 enzyme. We hypothesize that it could be an anchoring point for CaM in the full-length protein, facilitating activation of the kinase domain by an as yet unknown mechanism.

### NADKc-1 subcellular localization

Previous attempts to localize the plant CaM/Ca^2+^-dependent NAD^+^ kinase produced contradictory results. CaM-dependent NADK activity has been reported to be located in the non-plastidial cytosolic fraction (Simon et al., 1982), in the chloroplast (Jarrett et al., 1982), in the mitochondrial outer membrane (Sauer and Robinson, 1985) or in the chloroplast outer envelope membrane (Dieter and Marme, 1984). However, all these localizations may have been affected by fraction contamination.

Analysis of NADKc-1 topology, as predicted from its primary sequence by SPOCTOPUS (Viklund et al., 2008), indicated that the N-terminal region (NR) of the protein contains a stretch of amino acids (24 to 45) that may constitute a hydrophobic transmembrane helix (TM-helix). The C-terminus of the putative TM-helix contains a positively-charged lysine/arginine-rich sequence (Fig. S2), which is considered to be a hallmark of mitochondrial/chloroplast outer membrane proteins, or of proteins located in the endoplasmic reticulum (Lee et al., 2011). In addition, using the augmented Wimley-White scale (Jayasinghe et al., 2001), a maximum hydrophobicity of the TM-helix of0.28 was determined. Since this value is lower than 0.4, we hypothesize localization in the chloroplast/mitochondrial outer membrane (Lee et al., 2011).

To assess NADKc-1 localization *in vivo*, we produced a construct containing the NADKc-1 (N-terminal region - *NR*) fused to YFP, and another construct in which YFP was fused at the N-terminus of the whole NADKc-1 protein sequence. Both fusion proteins were inserted into plasmids under the control of the 35S promoter, and these constructs were used to transiently transform tobacco leaves via agrobacterium-mediated infiltration.

Confocal microscopy revealed the NADKc-1*NR*-YFP protein to be associated with small, fast moving organelles located in the periphery of epidermal cells. Co-infiltration with RFP fused with the first 69 amino acids of ATP synthase subunit to label mitochondria (Michaud et al., 2014) revealed co-localization of the YFP and RFP signals, thus suggesting that the NADKc-1 protein is associated with mitochondria (Fig. 2, top line). The NADKc-1*NR*-YFP signal also appeared more peripheral than the RFP signal, suggesting that the NADKc-1 protein might be associated with the outer mitochondrial membrane (Fig. 2, central lane), as predicted by our protein sequence analysis.

**Figure 2.**
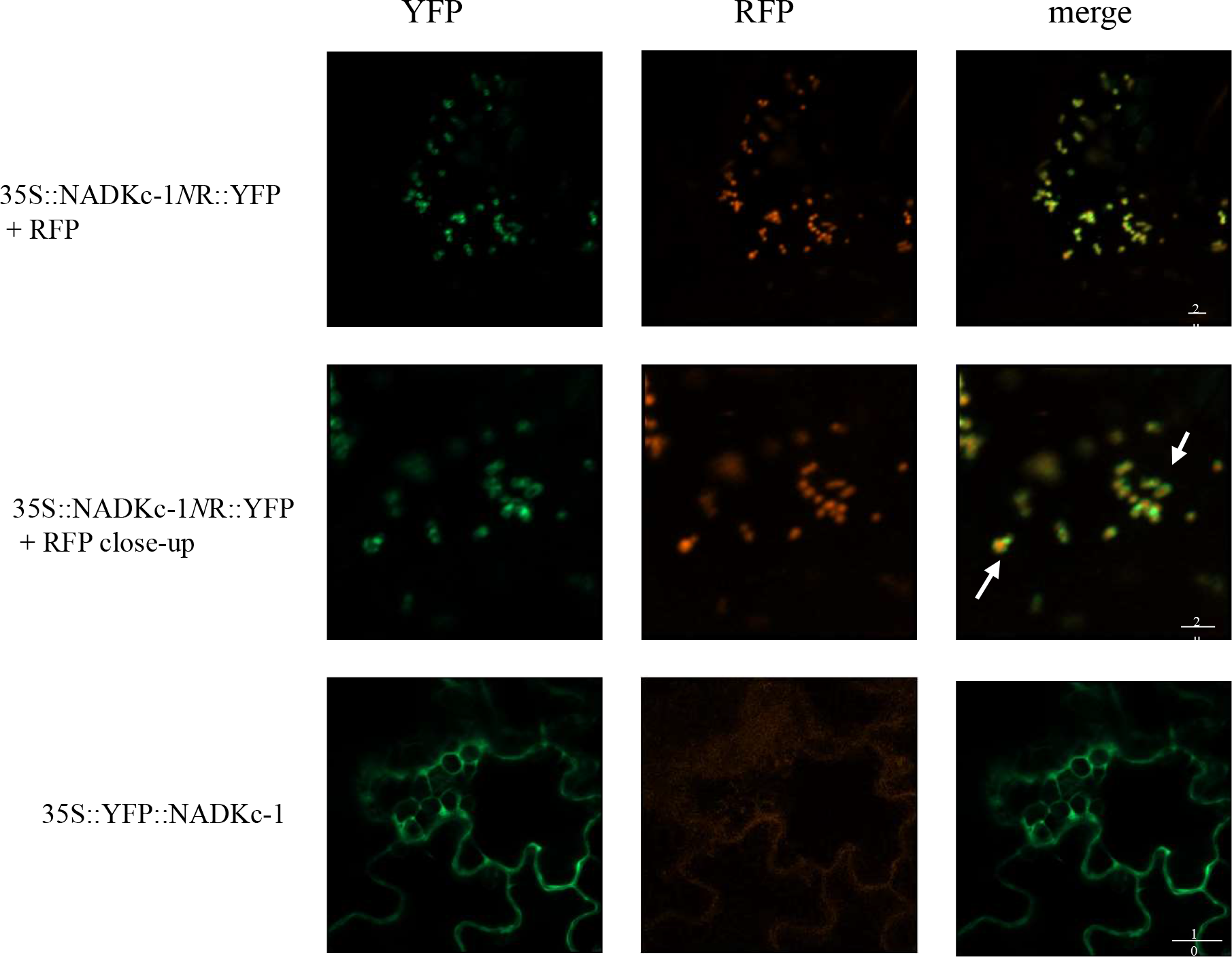
NADKc-1 associates with mitochondria in tobacco leaves. Representative pictures of transiently transformed tobacco leaf cells. Left: YFP (green), Center: RFP (red), and Right: merged fluorescence. The top row shows a representative image of tobacco cells co-transformed with the 35S::NADKc-1NR-YFP construct and a mitochondrial marker, pSu9– RFP. The middle row shows a close-up of mitochondria from the lower right region of the images in the top row. White arrows indicate regions in which the YFP signal appears peripheral with respect to the RFP signal. The bottom row shows a representative image of tobacco cells transformed with the 35S::YFP-NADKc-1 construct alone. The central image shows the background signal in the RFP channel.

The YFP-NADKc-1 construct, in which the YFP is linked to the N-ter of the whole NADKc-1 protein, was instead localized in the cytosol (Fig. 2, bottom line). This result was probably due to the NADKc-1 N-terminus being hidden in the middle of the sequence in this fusion protein, and corroborates the hypothesis that the *NR* of NADKc-1 is important for the protein to be correctly addressed to the mitochondria.

With few exceptions, plant CaMs and CaM-like proteins have mainly been observed in the cytosol (Abbas et al., 2014; Dell’ Aglio et al., 2016; Kushwaha et al., 2008; McCormack and Braam, 2003). The localization of NADKc-1 tethered to the outer mitochondrial membrane would allow interactions with these CaMs upon changes in the Ca^2+^ cytosolic concentration.

### NADKc-1 enhances Flg22 response in Arabidopsis seedlings

To investigate the physiological role of NADKc-1, we analyzed the two Arabidopsis T-DNA insertion lines SALK_006202 and GABI-KAT 311Hll, hereafter called *nadkc-1-1* and *nadkcl-2* (Fig. 3A). NADKc-1 transcripts were reduced by more than 95% in both lines (Fig. 3B).

**Figure 3:**
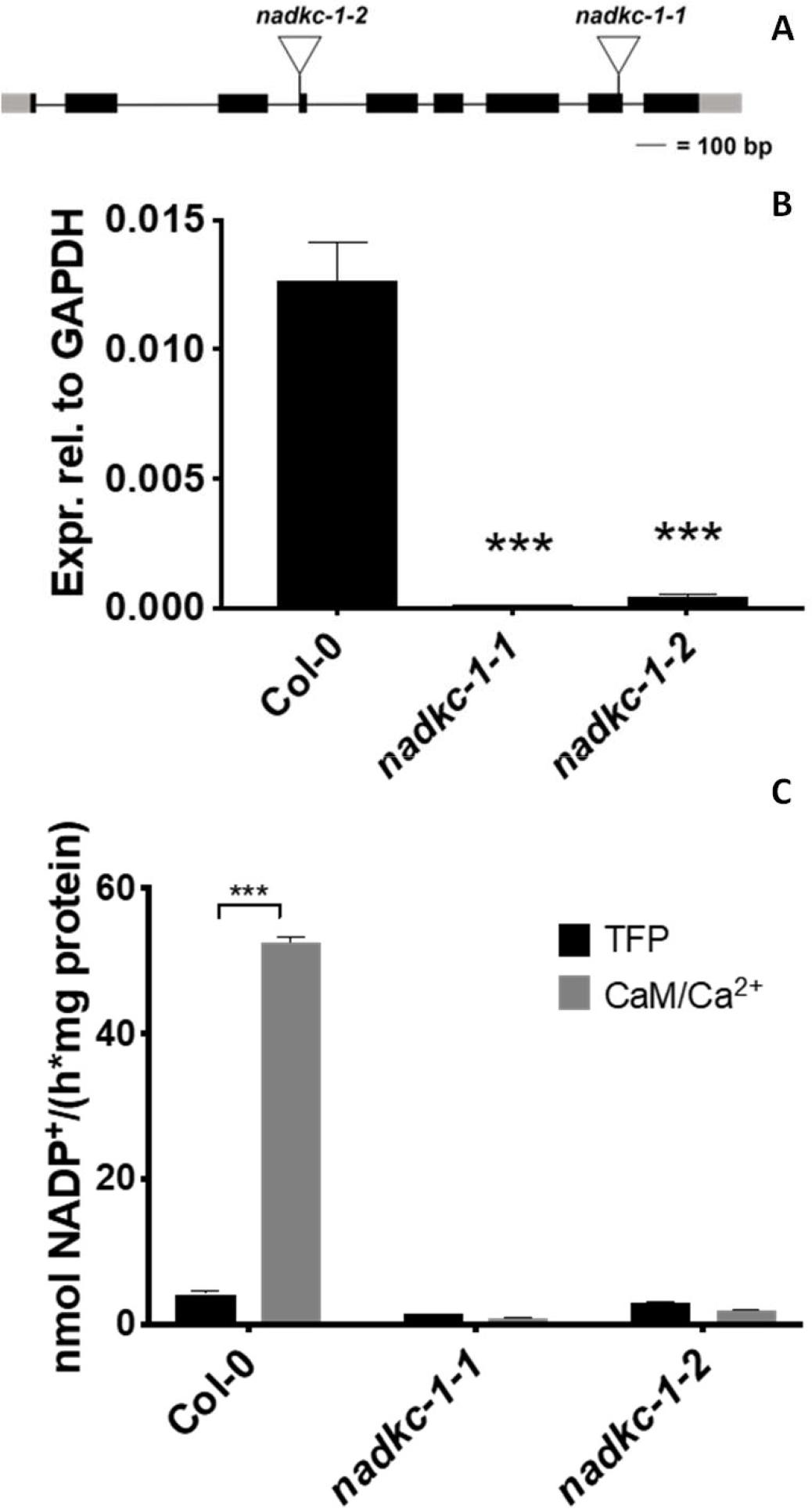
Characterization of Arabidopsis NADKc-1 mutant lines. **A:** schematic representation of the NADKc-1 gene and position of the T-DNA insertions in the *nadkc-1-1* and *nackc-1-2* mutant lines. **B:** *NADKc-1* transcript levels in Col-0 and NADKc-1-mutant seedlings. Levels are expressed relative to GAPDH. Data shown correspond to mean+/− s.d., n=3. **C:** NAD+ kinase activity measured in Col-0 and NADKc-1-mutant plants (7-day-old whole plantlets), in the presence of the CaM inhibitor TFP (40 μM) or AtCaM1 (250 nM) and Ca^2+^ (0.5 mM). Values correspond to the average of four replicates.

To confirm the unique role of the Atlg04280 gene product for CaM/Ca^2+^-dependent NADP^+^ production, NAD^+^ kinase activity was measured in protein extracts from WT and mutant seedlings, both in the presence of trifluoroperazine (TFP) - a CaM inhibitor - and CaM/Ca^2+^ (Fig. 3C). In Col-0 plants, the activity measured in the presence of AtCaM1/Ca^2+^ was more than 10-fold higher than in the presence of TFP, while in *nadkc-1-1/2* mutants, no difference was observed between the two conditions. In both mutants, activity levels were close to those measured in Col-0 plants in the presence of TFP, confirming the absence of NADKc-1 activity in these mutants.

Both *nadkc-1* mutants did not show any visible phenotype when grown under both short day and long day photoperiods (Fig. S5) and photosynthetic parameters (Fv/Fm, ETR and NPQ, (Maxwell and Johnson, 2000)) were the same in all genotypes (not shown). This suggested that the CaM-dependent NAD kinase is not involved in photosynthesis-driven growth.

As CaM-dependent NAD kinase activity was previously associated to the generation of the oxidative burst triggered by pathogen response (Grant et al., 2000; Harding et al., 1997), we expected to observe a decrease in the extracellular oxidative burst in *nadkc-1-1/2* mutants.

We therefore measured RBOH-dependent ROS production via a chemiluminescence assay on 7-day-old Arabidopsis seedlings. A dramatic reduction in ROS accumulation (more than 90%) was observed in the *nadkc-1-1/2* mutants (Fig. 4A). *Nadkc-1-1* seedlings stably transformed with the full-length protein under the control of the 35S promoter - *nadkc-1-l_NADKc-1-1* and *nadkc-1-l_NADKc-1-2* - had similar NADKc activity levels to the wild-type (Fig. 4B) and complemented the mutant phenotype by restoring physiological levels of ROS after flg22 challenge (Fig. 4C). These data support the idea that NADKc-1 plays a major role in triggering the pathogen MAMP-related oxidative burst.

**Figure 4:**
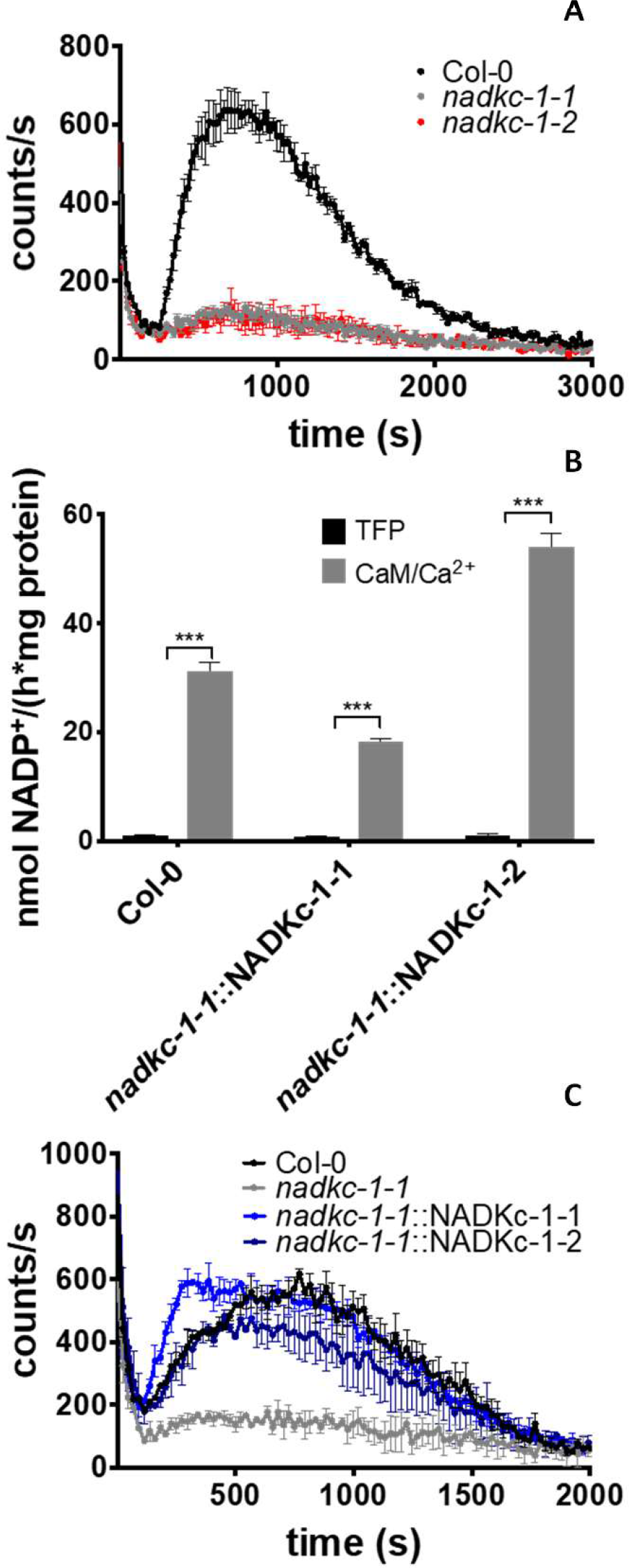
Flg22-induced extracellular oxidative burst requires NADKc-1 activity, which can be complemented in Arabidopsis mutant lines. **A:** Flg22 (1 μM)-induced oxidative burst in Col-0 and NADKc-1 mutant 7-day-old seedlings (30 plantlets per well, data shown correspond to mean+/− s.d. for 4 wells). **B.** NAD+ kinase activity measured in Col-0 and mutant plants complemented with NADKc-1 gene (nadkc-l_NADKc-1-1 and nadkc-l_NADKc-1-2) in 7-day-old whole plantlets, in the presence of the CaM inhibitor TFP (40 μM) or of AtCaM1 (250 nM) and Ca^2+^ (0.5 mM). **C.** Flg22 (1 μM)-induced oxidative burst in Col-0, *nadkc-1-1* mutant and mutant plants complemented with NADKc-1 gene (nadkc-l_NADKc-1-1 and nadkc-1_NADKc-1), 7-day-old seedlings, 30 plantlets per well, data shown correspond to mean +/− s.d. for 4 wells.

Based on these results, we propose that NADKc-1 is the protein responsible for CaM/Ca^2+^-dependent NAD^+^ kinase activity in Arabidopsis seedlings, and that it plays a role in biotic stress response as the most likely candidate providing NADP to the OPPP pathway that fuels production of ROS via RBOH.

### Distribution of CaM-dependent NAD kinase activity among plants and algae

A striking feature of NADKc-1 is the presence of a C-terminal kinase domain (amino acids 226-340) which was annotated as a “zeta toxin domain”. Modeling of the NADKc-1 C-terminus in Phyre 2 (Kelley et al., 2015) revealed that indeed this domain showed extensive similarity to the bacterial zeta toxin domain of the type II toxin-antitoxin systems (Yamaguchi et al., 2011). Type II zeta toxins-epsilon antitoxins systems are pairs of self-poisoning agents and their antidotes, which are common in Archaea and Bacteria, but rare in Eukaryotes (Yamaguchi et al., 2011). Possible roles of zeta toxins in plant virulence were recently revealed (Shidore and Triplett, 2017). Notably, the Xanthomonas AvrRxol zeta toxin is injected in plant cells during infections and is able to block the plant’s oxidative burst and its pathogen response (Shidore et al, 2017) by hijacking NAD metabolism via the production of the toxic compound 3’NADP.

The highest match for NADKc-1 in Phyre2 (99.7% confidence interval from amino acid 227 to amino acid 446) was observed for the *Streptococcus pneumoniae* zeta toxin PezT protein (PDB code: 2P5T, (Khoo et al., 2007; Mutschler et al., 2011), Supplementary Fig. S6A and B). In contrast, we could not find structural homologs for the N-terminal domain.

Mining of public databases indicated that the domain organization of NADKc-1 (*i.e.*, a ca 200 amino acid domain of unknown function at the N-terminus with a putative CBS followed by a kinase domain with a Walker Motif annotated “zeta toxin domain”) was only found in higher plants and some alga (see below) among eukaryotes.

To better trace the evolution of the plant CaM-dependent NAD kinase, we compared gene sequences and NADK activity in several plants and algae.

Firstly, we built a maximum likelihood phylogenetic tree with representative putative NADKc proteins of plants and algae (Fig. S6C). The phylogenetic tree showed that plant NADKc-like proteins form four major clusters that correspond to the main plant phylogenetic groups with the exception of Gymnosperms and Pteridophytes, in which the NADKc gene has apparently been lost. Many plants, especially Dicots, contain several genes encoding for this protein, suggesting duplication events across evolution.

We also looked at NADKc-like proteins in the genomes of available annotated algae. While the genomes of *Chlamydomonas reinhardtii*, *Ostreococcus taurii* and *Chara braunii* appeared devoided of NADKc-like sequences, the genomes of *Coccomyxa subellipsoidea*, *Ulva mutabilis*, *Klebsormidiumflaccidum*, *Spyrotaenia minuta*, *Entransiafimbriata*, Mougeotia sp. and Spirogyra sp. all harbor one NADK-like sequence with a conserved zeta toxin domain and a long N-terminal domain with unknown function. Among the latter species, genes of *Klebsormidiumjlaccidum*, *Entransia fimbriata*, Mougeotia sp. and Spirogyra sp. contain a clear CBS, while in *Coccomyxa subellipsoidea*, *Ulva mutabilis* and *Spirotaenia minuta*, this region is less conserved (Fig S3 and S6). In agreement with the genomic survey, CaM-dependent NAD kinase activity could be successfully measured in the moss *M. polymorpha* and filamentous alga *K. flaccidum* but not in the unicellular alga *C. reinhardtii* (Supplemental Table S3). Interestingly, both the total CaM-dependent NAD kinase activity and the percentage of CaM-dependent NADK-activity on the total NADK activity increase from *K. jlaccidum* (4.0 nmol/h/mg; 66.7%) to *M. polymorpha* (5.9 nmol/h/mg; 85.7%) and *to A. thaliana* (30.2 nmol/h/mg; 96.8%). It is therefore possible that the importance of the CaM control on NADKc evolved in concomitance during plant colonization of land thereby becoming a key element in plant response to pathogen elicitors. Overall, our data allows proposing that the CaM-dependent NAD kinase activity is only found in photosynthetic species carrying NADKc-1 related proteins, which would represent the only proteins harboring CaM-dependent NAD kinase activity in plants and algae.

## METHODS

### Chemicals

All chemicals were from Sigma Aldrich.

### Plant growth and isolation of homozygous NADKc-1 lines

*A. thaliana* Col-0 ecotype was used in this study. Plants were grown under 65% humidity and either long day (16 h light-85 μE, 8 h dark) or short day (8 h light-I 90 μE, 16 h dark) conditions. Day-time temperature was set to 20 °C, and night-time temperature to 18 °C.

The two T-DNA insertion lines, *nadkcl-1* (SALK_006202) and *nadkcl-2* (GABI_311Hl1), were obtained from NASC/ABRC(Alonso et al., 2003; Kleinboelting et al., 2012). Lines were selected in the appropriate antibiotic(kanamycin for *nadkcl-1* and sulfadiazine for *nadkcl-2*) and genotyped by PCR using left border primers LBbl.3 *(nadkcl-1)* or LB GABI-KAT *(nadkcl-2)* and the appropriate specific primers listed in Supplemental Table S4. PCR products were sequenced to confirm the precise position of each insertion.

*Klebsormidium jlaccidum* (Hori et al., 2014) (SAG335-2b curated as *Klebsormidium nitens)* was obtained from EPSAG (Department of Experimental Phycology and Culture Collection of Algae, Gottingen Universitat, Germany). The alga was grown on agar plates under continuous light(60 μE) in the Modified Bolds 3N Medium (https://utex.org/products/modified-bolds-3nmedium) without vitamins.

*M. polymorphawas* collected in the forest(GPS coordinate: 45.335088, 5.632257) and *Chlamydomonas reinhardtii* (C137 strain) was grown in TAP medium at 24°C under continuous mild white light (40 μmol photons m^−2^ s^−1^) exposure. Protein extracts were prepared as described in the supplementary information for Arabidopsis.

Additional Material and Methods procedures are described in the Supplementary Information.

### Accession numbers

Sequence data from this article can be found in the EMBL/GenBank data libraries under accession number At1g04280; UniProt accession QOWUYl.

