## Supplemental information for "Identification of the Calmodulin-dependent NAD^+^ kinase sustaining the elicitor-induced oxidative burst in plants"

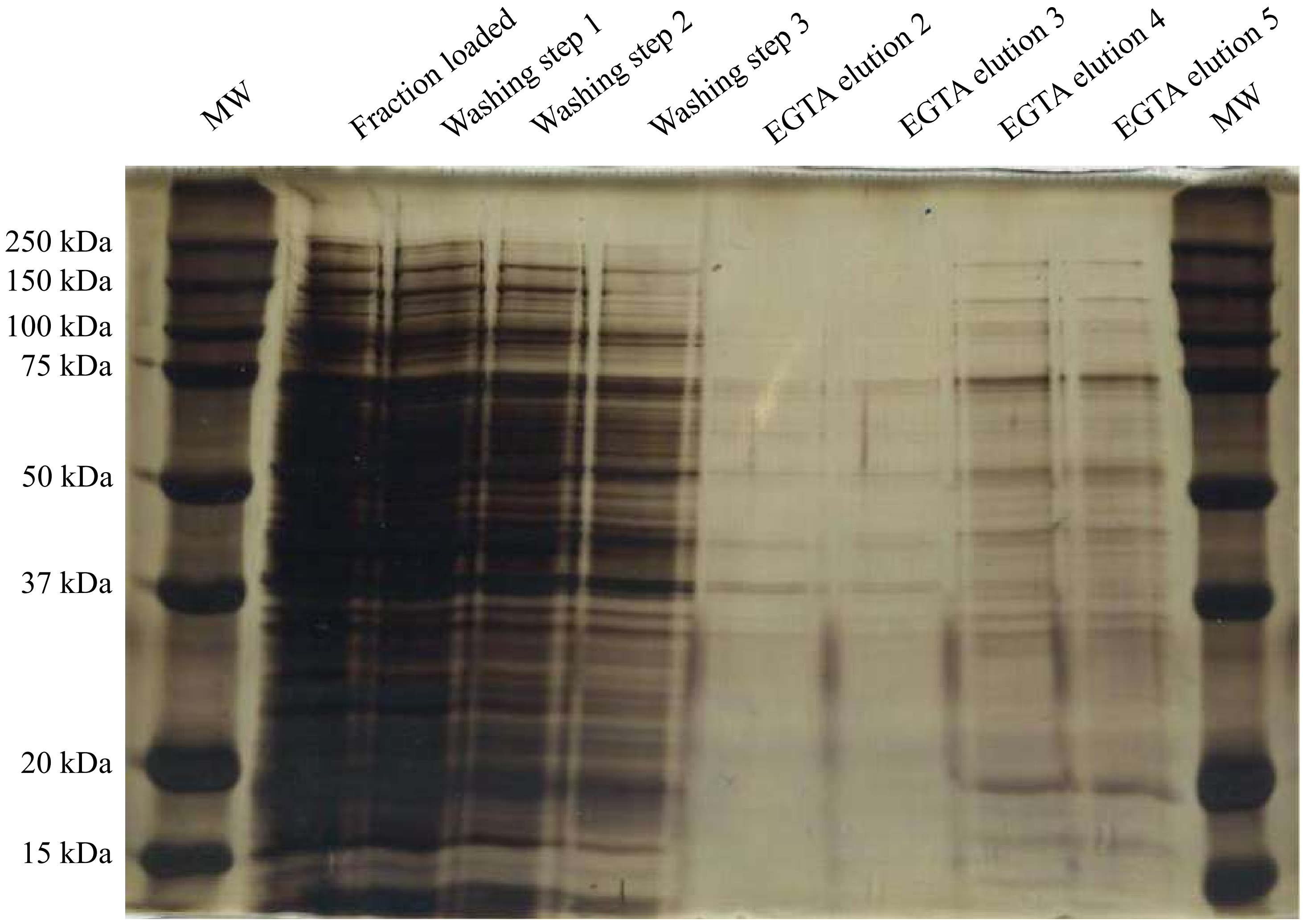


**Supplementary Figure S1: CaM-affinity purification of native NADKc from Arabidopsis plantlets.** Proteins loaded and eluted from the CaM affinity chromatography column were separated by SDS-PAGE and stained with silver nitrate. Mass spectrometry-based proteomics was used to identify proteins strongly enriched in the EGTA elution compared to the $Ca^{2+}$-containing washing steps (Supplemental Table S1).

MVKPLGEGTRDVSNVGCKSVDLLLASLGGLAAAVAAAYAGELLLRRRKLDQGASMGYKDV
KIAPLIERKDSGRRSNLERFSHYVARQLGFEDPNEYPQLCKLANGYLLKTKGYDENVDEY
LENEAERDSLYVHLLEEFDRCILTYFSFNWTQSSNLISQALSDESDQKVPKLKDFVMAAT
RKQRFERVTKDLKVKRVISTLVEEMRVIGSGSSEPHCTEVMSPVAHNKRSPVLLLMGGGM
GAGKSTVLKDIFLESFWSEAQADAVVIEADAFKETDVIYRALSSRGHHDDMLQTAELVHQ
SSTDAASSLLVTALNDGRDVIMDGTLSWEPFVEQMIEMARNVHKQKYRMGEGYKVSEEGT
ITEKYWEEEEEETKENGKQQNLKPYRIELVGVVCDAYLAVARGIRRALMVKRAVRVKPQL
NSHKRFANAFPKYCELVDNARLYCTNAVGGPPRLIAWKDGNSKLLVDPEDIDCLKRVSSL
NPDAESIYELYPDPSQLSKPGSVWNDVVLVPSRPKVQKELSDAIRRIEKAQPKN

**Supplementary Fig. S2. SPOCTOPUS-predicted topology**. SPOCTOPUS result was generated from http://octopus.cbr.su.se/ using the full-length predicted amino acid sequence for NADKc1. The underlined sequence corresponds to predicted transmembrane-helix residues; residues represented in red correspond to the lysine/arginine-rich region.

### Walker A motif — A

| | |
|---|---|
| *A. thaliana-1* (At1g04280) | MGGGMGAGKSTVLKD |
| **Plants** | |
| *S. phallax-1* | IGGGMGAGKSTVVQE |
| *P. patens* | IGGGMGAGKSTIVKE |
| *M. polymorpha* | IGGGMGAGKSTVVKE |
| *S. mollendorfii-1* | MGGGMGAGKSTVLKD |
| *S. lycopersicum1* | MGGGMGAGKSTVLKD |
| *G. max-1* | MGGGMGAGKSTVLKD |
| *M. esculenta-1* | MGGGMGAGKSTVIKD |
| *P. trichocarpa-1* | MGGGMGAGKSTVTKD |
| *M. acuminata-1* | MGGGMGAGKSTVLKE |
| *O. sativa-1* | MGGGMGAGKSTVLKE |
| *B. distachyon-1* | MGGGMGAGKSTVLKE |
| *S. italica-1* | MGGGMGAGKSTVLME |
| *Z. mays-1* | MGGGMGAGKSTVLKE |
| **Algae** | |
| *C. Subellipsoidea* | LGGGMAAGKSTVREI |
| *U. mutabilis* | LGGGMAAGKSSVRNE |
| *S. minuta* | LGGGMGAGKSTAVKE |
| *K. flaccidum* | MAGGMGAGKSTVRQE |
| *E. fimbriata* | MGGGMGAGKSTAARR |
| *Mougeotia sp.* | IGGGMGAGKSTV-KE |
| *Spirogyra sp.* | LGGGMGAGKSTVAKQ |

### CaM binding site — B

| | |
|---|---|
| *A. thaliana-1* (At1g04280) | KVPKLKDFVMAATRKQRFERVTKDLKVKR |
| **Plants** | |
| *S. phallax-1* | -QKRLRKAVWLATRPQRIERVLKCLKTKR |
| *P. patens* | -EKKLHKAVLCALKKQRYEAVLKSLSTKR |
| *M. polymorpha* | QKRSLKKAVLSATRKQRYQRVMQDLKTKR |
| *S. mollendorfii-1* | ---NLRSIVMAATRKHRFQRAMQTLKAKR |
| *S. lycopersicum-1* | -QKRLKDLVLAATRKQRFEKITKDLKVTR |
| *G. max-1* | QKKKLKGILLAATREQRFDRVTKNLKVTR |
| *M. esculenta-1* | KRHKLKDVVLAATRKQRFERVNKELKVTR |
| *P. trichocarpa-1* | KKPKLKGIVMAATRKQRFERVTKNLKVTR |
| *M. acuminata-1* | QKMKLKNFVMEATRKLRFERVTKDLKVTR |
| *O. sativa-1* | SKKKLRNLVLEATRKQRFERVTRDLKVTR |
| *B. distachyon-1* | TRKKLRNLVLEATRKQRFERVTRDLKVTR |
| *S. italica-1* | STKKLRNVFMEATRKQRFARVTRDLKVTR |
| *Z. mays-1* | SKRKLRNMVLEATRKQRFERVTRDLKVTR |
| **Algae** | |
| *K. flaccidum* | KRHSLKRAVLSATRTQRYKQLLRSLGPQR |
| *E. fimbriata* | -K—SLKGAVLRATRTHRYERLVTALGSQR |
| *Mougeotia sp.* | KKMTFKAAVRKATSIQRVKKVIEHLGPQR |
| *Spirogyra sp.* | RL-SFKKAVLNATRNQRMEKIMTNFKTQQ |

**Supplementary Fig. S3: Features of NADKc-1 primary sequence and its homologues in other plant and algae species. A)** Walker A motif (in red, residues characterizing the motif; *A. thaliana* NADKc-1 amino acids 236-250); **B)** putative CaM binding site (*A. thaliana* NADKc-1 amino acids 168-200). Red residues are those constituting the 1-5-8-14 motif. This domain is not conserved in *C. Subellipsoidea, U. Mutabilis and S. minuta.*

*A. thaliana*: *Arabidopsis thaliana*, *S. phallax*: *Sphagnum phallax*, *P. patens*: *Physcomitrella patens*, *M. polymorpha*: *Marchantia polymorpha*, *S. mollendorffii*: *Selaginella mollendorffii*, *S. lycopersicum*: *Solanum lycopersicum*, *G. max*: *Glycine max*, *M. truncatula*: *Medicago truncatula*, *P. trichocarpa*: *Populus trichocarpa*, *M. acuminata*: *Musa acuminata*, *O. sativa*: *Oryza sativa*, *B. distachyon*: *Brachypodium distachyon*, *S. italica*: *Setaria italica*, *Z. mays*: *Zea mays*, *C. subellipsoidea*: *Coccomyxa subellipsoidea*, *U. mutabilis*: *Ulva mutabilis*, *S. minuta*: *Spirotaenia minuta*, *K. flaccidum*: *Klebsormidium flaccidum*, *E. fimbriata*: *Entransia fimbriata*. For gene references, see the legend of Fig. S7.

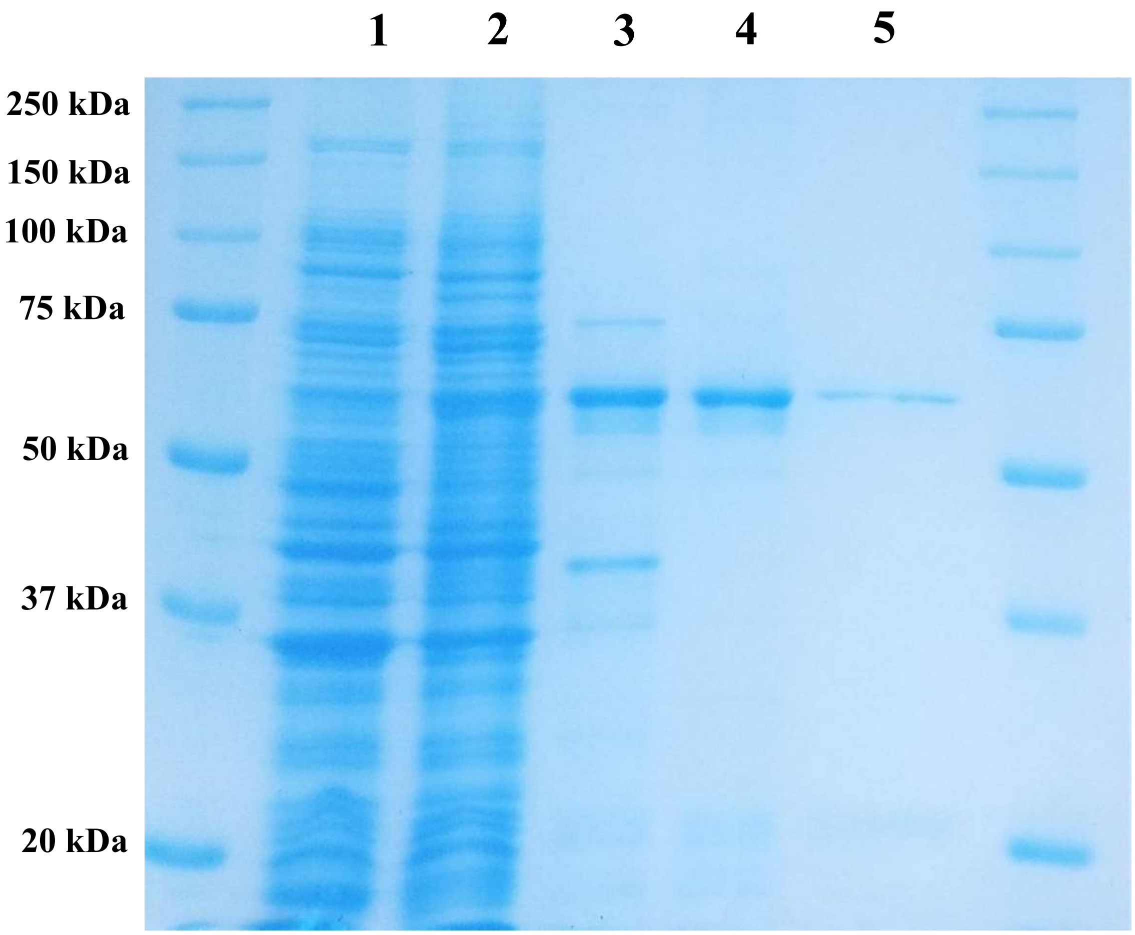


**Supplementary Fig. S4: Purification of recombinant NADKc-1 produced in *E. coli*.** 6HIS-D38NADKc-1 was purified as indicated in Material and Methods. Lane 1: Rosetta$^2$ soluble extract transformed with empty pET28a (40 µg); lane 2: Rosetta$^2$ soluble extract transformed with pET28a-6HIS-Δ38NADKc-1 (40 µg); lane 3: Ni-NTA pool (10 µg); lane 4: urea-denatured 6HIS-Δ38NADKc-1 purified on Ni-NTA (10 µg); lane 5: refolded 6HIS-Δ38NADKc-1 (2 µg).

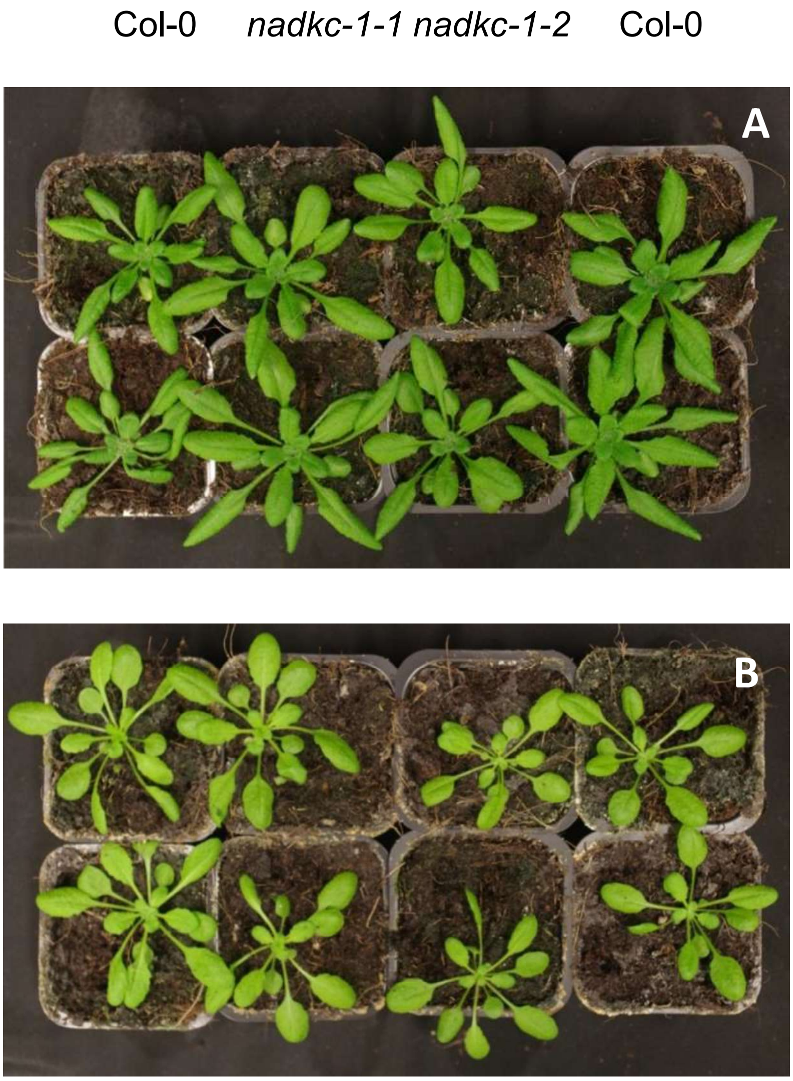


**Supplementary Fig. S5: Phenotype of *nadkc-1* mutants. A)** 21-day-old plants grown under long day photoperiod (16 h light/8 h dark); **B)** 28-day-old plants grown under short day photoperiod (8 h light/16 h dark).

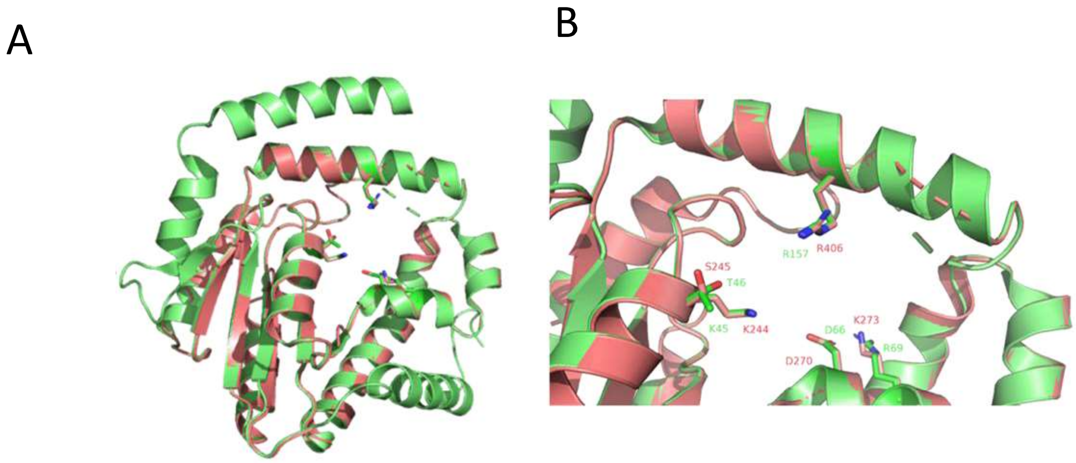


**Supplementary Figure S6: Structural model of NADKc-1. A:** superposition of the NADKc-1 structural model generated with Phyre2 (red) and the PezT protein from *S. pneumoniae* (green, PDB ID: 2P5T). **B:** close-up on the putative ATP-binding site, with conserved ATP coordinating amino-acids highlighted.

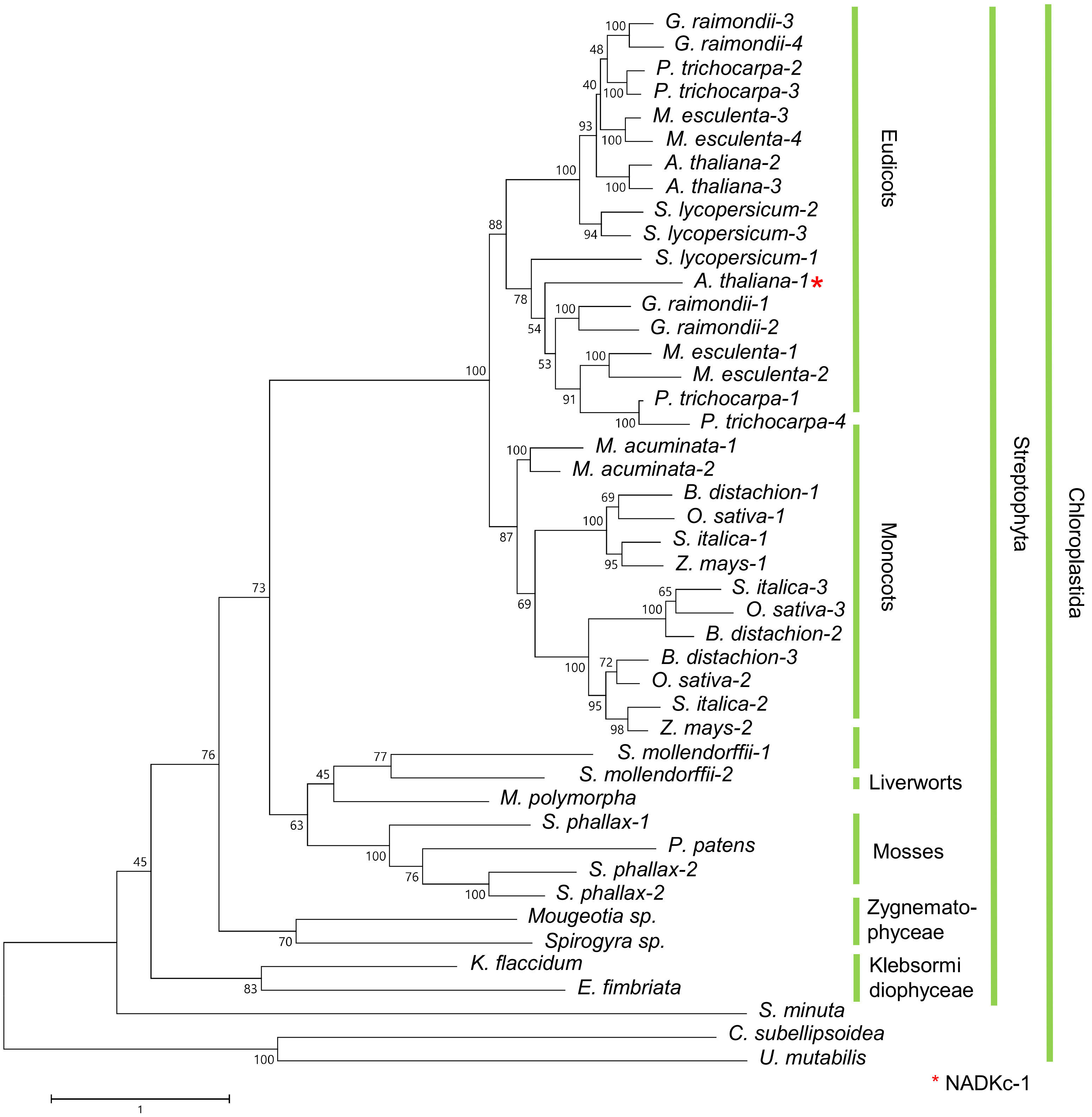


**Supplementary Figure S7: Phylogeny of NADKc-1.** Maximum likelihood phylogenetic tree of NADKc-like protein in the following plant and algal species : *Arabidopsis thaliana* (*A. thaliana*), *Populus trichocarpa* (*P. trichocarpa*), *Manihot esculenta* (*M. esculenta*), *Gossypium raimondii* (*G. raimondii*), *Solanum lycopersicum* (*S. lycopersicum*), *Setaria italica* (*S. italica*), *Zea mays* (*Z. mays*), *Oryza sativa* (*O. sativa*), *Brachypodium distachyon* (*B. distachyon*), *Musa acuminata* (*M. acuminata*), *Marchantia polymorpha* (*M. polymorpha*), *Physcomitrella patens* (*P. patens*), *Sphagnum phallax* (*S. phallax*), *Selaginella mollendorffii* (*S. mollendorffii*) *Klebsormidium flaccidum* (*K. flaccidium*), *Coccomyxa subellipsoidea* (*C. subellipsoidea*), *Ulva mutabilis* (*U. mutabilis*), *Spirotaenia minuta* (*S. minuta*), Mougeotia sp., Spirogyra sp. and *Entransia fimbriata* (*E. fimbriata*). Accession numbers in the next slide…/…

…/… Accession numbers: *A. thaliana-1* (NADKc-1): At1g04280, *A. thaliana-2*: At1g06750, *A. thaliana-3*: At2g30630, *P. trichocarpa-1*: Potri.008G162000, *P. trichocarpa-2*: Potri.002G043000, *P. trichocarpa-3*: Potri.005G220000, *P. trichocarpa-4*: Potri.008G162400, *M. esculenta-1*: Manes.15G036900, *M. esculenta-2*: Manes.03G168900, *M. esculenta-3*: Manes.01G200600, *M. esculenta-4*: Manes.05G086200, *G. raimondii-1*: Gorai.011G171100, *G. raimondii-2*: Gorai.006G246600, *G. raimondii-3*: Gorai.004G074100, *G. raimondii-4*: Gorai.013G104300, *S. lycopersicum-1*: Solyc06g053810, *S. lycopersicum-2*: Solyc06g031670, *S. lycopersicum-3*: Solyc08g059750, *S. italica-1*: Seita.9G163700, *S. italica-2*: Seita.3G185100, *S. italica-3*: Seita.5G337000, *Z. mays-1*: GRMZM2G070252, *Z. mays-2*: GRMZM2G368410, *O. sativa-1*: Os03g43010, *O. sativa-2*: Os05g43300, *O. sativa-3*: Os01g56764, *B. distachyon-1*: Bradi1g14307, *B. distachyon-2*: Bradi2g51490, *B. distachyon-3*: Bradi2g20400, *M. acuminata-1*: SMUA_Achr5T03210, *M. acuminata-2*: GSMUA_Achr7T01560, *M. polymorpha*: Mapoly0142s0012, *P. patens*: Pp3c2_3490V3, *S. phallax-1*: Sphallax0059s0037, *S. phallax-2*: Sphallax0011s0002, *S. phallax-3*: Sphallax0120s0019, *S. mollendorffii-1*: scaffold 73427, *S. mollendorfii-2*: scaffold 231175, *K. flaccidum*: kfl00274_0130, *C. subellipsoidea*: XP_005648203, *U. mutabilis*: UM028_0076.1, *S. minuta*: NNHQ_2000691, *Mougeotia*: ZRMT_2002068, *Spirogyra*: HAOX_2025158, *E. fimbriata*: BFIK_2025349. The tree was generated with ClustalW and MEGA.

**Table S1. Mass spectrometry-based proteomics identification of proteins strongly enriched in the EGTA elution compared to the Ca2+-containing washing steps of the CaM affinity chromatography.**

Only proteins identified only in Eluate sample with SSC ≥ 5 or enriched at least 10 times in Eluate sample compared to Flowthrough sample are shown. In orange: NADKc-1.

Pep: Number of identified peptides

SC: Spectral Counts: mass spectra allow the identification of peptides belonging to the protein

SSC: Spectral Counts: mass spectra allow the identification of peptides belonging to the protein

| Accession | Description | Gene name | Coverage | Mol. Weight | SSC Eluate / SSC Flowthrough | Flowthrough | | | Eluate | | |
|---|---|---|---|---|---|---|---|---|---|---|---|
| | | | | | | Pep | SC | SSC | Pep | SC | SSC |
| RNJ_ARATH | Ribonuclease J | RNJ | 33,37 | 100554 | Eluate cnly | | | | 29 | 34 | 34 |
| RH42_ARATH | DEAD-box ATP-dependent RNA helicase 42 | RH42 | 20,24 | 133033 | Eluate cnly | | | | 25 | 28 | 28 |
| ICR1_ARATH | Interactor of constitutive active ROPs 1 | ICR1 | 52,03 | 38358 | Eluate cnly | | | | 19 | 22 | 22 |
| PARNL_ARATH | Poly(A)-specific ribonuclease PARN-like | At3g25430 | 24,43 | 68807 | Eluate cnly | | | | 16 | 18 | 18 |
| A0A178W9E9_ARATH | Obg-like ATPase 1 | AXX17_At1g50720 | 28,27 | 45628 | Eluate cnly | | | | 10 | 17 | 17 |
| MED16_ARATH | Mediator of RNA polymerase II transcription subunit 16 | MED16 | 15,81 | 138187 | Eluate cnly | | | | 15 | 17 | 17 |
| Q9SYM7_ARATH | 2-oxoglutarate (2OG) and Fe(II)-dependent oxygenase superfamily protein | At1g78550 | 26,97 | 41015 | Eluate cnly | | | | 10 | 14 | 14 |
| TON1A_ARATH | Protein TONNEAU 1a | TON1A | 37,31 | 29334 | Eluate cnly | | | | 13 | 14 | 13 |
| ODP23_ARATH | Dihydrolipoyllysine-residue acetyltransferase component 3 of pyruvate dehydrogenase complex, mitochondrial | At1g54220 | 23,19 | 58467 | Eluate cnly | | | | 11 | 14 | 13 |
| PHOT2_ARATH | Phototropin-2 | PHOT2 | 15,08 | 102472 | Eluate cnly | | | | 15 | 15 | 13 |
| STA1_ARATH | Protein STABILIZED1 | STA1 | 11,66 | 115576 | Eluate cnly | | | | 12 | 13 | 13 |
| UGPA2_ARATH | UTP--glucose-1-phosphate uridylyltransferase 2 | UGP2 | 25,59 | 51738 | Eluate cnly | | | | 10 | 13 | 13 |
| ATPG1_ARATH | ATP synthase gamma chain 1, chloroplastic | ATPC1 | 31,37 | 40911 | Eluate cnly | | | | 10 | 12 | 12 |
| EBFC2_ARATH | Nucleoid-associated protein At2g24020, chloroplastic | STIC2 | 44,51 | 19812 | Eluate cnly | | | | 13 | 15 | 12 |
| PR35B_ARATH | Pre-mRNA-processing protein 40B | PRP40B | 15,93 | 113569 | Eluate cnly | | | | 11 | 12 | 12 |
| PSD3_ARATH | Phosphatidylserine decarboxylase proenzyme 3 | PSD3 | 17,32 | 70352 | Eluate cnly | | | | 10 | 12 | 12 |
| SUOX_ARATH | Sulfite oxidase | SOX | 34,1 | 43329 | Eluate cnly | | | | 10 | 12 | 12 |
| DOT2_ARATH | SART-1 family protein DOT2 | DOT2 | 17,32 | 94142 | Eluate cnly | | | | 10 | 11 | 11 |
| COIL_ARATH | Coilin | COIL | 12,5 | 68667 | Eluate cnly | | | | 9 | 11 | 11 |
| NEDD1_ARATH | Protein NEDD1 | NEDD1 | 20,97 | 84757 | Eluate cnly | | | | 9 | 11 | 11 |
| PTA16_ARATH | Protein PLASTID TRANSCRIPTIONALLY ACTIVE 16, chloroplastic | PTAC16 | 26,47 | 54358 | Eluate cnly | | | | 9 | 10 | 10 |
| A0A178V7A8_ARATH | Uncharacterized protein | AXX17_At4g18270 | 27,95 | 43910 | Eluate cnly | | | | 8 | 10 | 10 |
| E135_ARATH | Glucan endo-1,3-beta-glucosidase 5 | At4g31140 | 21,28 | 52715 | Eluate cnly | | | | 8 | 10 | 10 |
| SH3P3_ARATH | SH3 domain-containing protein 3 | SH3P3 | 28,77 | 39534 | Eluate cnly | | | | 8 | 10 | 10 |
| TON1B_ARATH | Protein TONNEAU 1b | TON1B | 30,74 | 29185 | Eluate cnly | | | | 10 | 10 | 9 |
| PDPK1_ARATH | 3-phosphoinositide-dependent protein kinase 1 | PDPK1 | 18,74 | 54711 | Eluate cnly | | | | 8 | 9 | 9 |
| A0A178USK6_ARATH | Uncharacterized protein | AXX17_At4g37900 | 23,19 | 56571 | Eluate cnly | | | | 9 | 9 | 8 |
| Q0WUY1_ARATH | p-loop containing nucleoside triphosphate hydrolases superfamily protein | At1g04280 | 15,33 | 60166 | Eluate cnly | | | | 8 | 8 | 8 |
| A0A1P8BE11_ARATH | RabGAP/TBC domain-containing protein | At5g52580 | 19,15 | 78317 | Eluate cnly | | | | 7 | 8 | 8 |
| AT13A_ARATH | Autophagy-related protein 13a | ATG13A | 11,44 | 66564 | Eluate cnly | | | | 6 | 8 | 8 |
| C3H38_ARATH | Zinc finger CCCH domain-containing protein 38 | At3g18640 | 11,24 | 75581 | Eluate cnly | | | | 7 | 8 | 8 |
| NFXL1_ARATH | NF-X1-type zinc finger protein NFXL1 | NFXL1 | 8,92 | 130716 | Eluate cnly | | | | 8 | 8 | 8 |
| PP1R8_ARATH | Protein phosphatase 1 regulatory inhibitor subunit PPP1R8 homolog | At5g47790 | 16,26 | 39985 | Eluate cnly | | | | 5 | 8 | 8 |
| PUX9_ARATH | Plant UBX domain-containing protein 9 | PUX9 | 23,24 | 52136 | Eluate cnly | | | | 7 | 8 | 8 |
| TPLAT_ARATH | Protein TPLATE | TPLATE | 11,39 | 130908 | Eluate cnly | | | | 8 | 8 | 8 |
| A0A178UB82_ARATH | Uncharacterized protein | AXX17_At5g20380 | 16,95 | 39678 | Eluate cnly | | | | 7 | 7 | 7 |
| A0A178V0X9_ARATH | Uncharacterized protein | AXX17_At4g45060 | 13,86 | 69578 | Eluate cnly | | | | 7 | 7 | 7 |
| A0A178V6Z0_ARATH | Uncharacterized protein | AXX17_At4g35270 | 12,79 | 65126 | Eluate cnly | | | | 6 | 7 | 7 |
| A0A178VQ14_ARATH | PIA1 | AXX17_At2g16010 | 25,81 | 30497 | Eluate cnly | | | | 6 | 7 | 7 |
| A0A178W283_ARATH | Uncharacterized protein | AXX17_At2g15630 | 12,94 | 71331 | Eluate cnly | | | | 5 | 7 | 7 |
| A0A178WAH4_ARATH | Uncharacterized protein | AXX17_At1g67070 | 26,75 | 40674 | Eluate cnly | | | | 7 | 7 | 7 |
| BIM2_ARATH | Transcription factor BIM2 | BIM2 | 32,15 | 34487 | Eluate cnly | | | | 7 | 7 | 7 |
| DNLI1_ARATH | DNA ligase 1 | LIG1 | 11,14 | 87740 | Eluate cnly | | | | 7 | 7 | 7 |
| F4K465_ARATH | Nucleoporin-like protein | At5g20200 | 12,86 | 81685 | Eluate cnly | | | | 7 | 7 | 7 |
| GEML1_ARATH | GEM-like protein 1 | FIP1 | 26,64 | 27958 | Eluate cnly | | | | 6 | 7 | 7 |
| PTN2B_ARATH | Phosphatidylinositol 3,4,5-trisphosphate 3-phosphatase and protein-tyrosine-phosphatase PTEN2B | PTEN2B | 19,94 | 70073 | Eluate only | | | | 7 | 7 | 7 |
| TAF4B_ARATH | Transcription initiation factor TFIID subunit 4b | TAF4B | 11,03 | 93741 | Eluate only | | | | 6 | 7 | 7 |
| TYW23_ARATH | tRNA wybutosine-synthesizing protein 2/3/4 | At4g04670 | 6,23 | 110856 | Eluate only | | | | 5 | 7 | 7 |
| A0A178UKW5_ARATH | Uncharacterized protein | AXX17_At5g61520 | 16,78 | 50341 | Eluate only | | | | 6 | 6 | 6 |
| A0A178UGE6_ARATH | Uncharacterized protein | AXX17_At5g58590 | 20,97 | 48903 | Eluate only | | | | 6 | 6 | 6 |
| A0A178UX40_ARATH | Uncharacterized protein | AXX17_At4g39230 | 33,33 | 15999 | Eluate only | | | | 8 | 12 | 6 |
| A0A178W3D1_ARATH | Uncharacterized protein | AXX17_At1g75860 | 29,11 | 24258 | Eluate only | | | | 4 | 6 | 6 |
| A0A178WJI9_ARATH | Uncharacterized protein | AXX17_At1g72510 | 19,93 | 31907 | Eluate only | | | | 6 | 6 | 6 |
| A0A178WC39_ARATH | Uncharacterized protein | AXX17_At1g53280 | 9,87 | 86016 | Eluate only | | | | 6 | 6 | 6 |
| A0A178WE72_ARATH | Uncharacterized protein | AXX17_At1g25550 | 4,15 | 166231 | Eluate only | | | | 5 | 6 | 6 |
| DUF7_ARATH | DUF724 domain-containing protein 7 | DUF7 | 9,97 | 80740 | Eluate only | | | | 6 | 6 | 6 |
| PATL3_ARATH | Patellin-3 | PATL3 | 10,61 | 56105 | Eluate only | | | | 4 | 6 | 6 |
| PDI21_ARATH | Protein disulfide-isomerase like 2-1 | PDIL2-1 | 18,84 | 39497 | Eluate only | | | | 6 | 6 | 6 |
| SLU7A_ARATH | Pre-mRNA-splicing factor SLU7-A | At1g65660 | 10,47 | 61978 | Eluate only | | | | 5 | 6 | 6 |
| SUR1_ARATH | S-alkyl-thiohydroximate lyase SUR1 | SUR1 | 17,97 | 51088 | Eluate only | | | | 6 | 6 | 6 |
| ICR4_ARATH | Interactor of constitutive active ROPs 4 | ICR4 | 24,38 | 36057 | Eluate only | | | | 4 | 5 | 5 |
| A0A178VKF6_ARATH | Uncharacterized protein | AXX17_At3g12680 | 6,65 | 73124 | Eluate only | | | | 4 | 5 | 5 |
| A0A178W2B9_ARATH | Uncharacterized protein | AXX17_At1g09880 | 14,78 | 54855 | Eluate only | | | | 5 | 5 | 5 |
| AT18H_ARATH | Autophagy-related protein 18h | ATG18H | 5,72 | 100514 | Eluate only | | | | 5 | 5 | 5 |
| BLI_ARATH | Protein BLISTER | BLI | 7 | 78384 | Eluate only | | | | 5 | 5 | 5 |
| CBSX2_ARATH | CBS domain-containing protein CBSX2, chloroplastic | CBSX2 | 16,81 | 25956 | Eluate only | | | | 3 | 5 | 5 |
| DEG15_ARATH | Glyoxysomal processing protease, glyoxysomal | DEG15 | 6,63 | 76126 | Eluate only | | | | 4 | 5 | 5 |
| HIS4_ARATH | Imidazole glycerol phosphate synthase hisHF, chloroplastic | HISN4 | 11,66 | 64192 | Eluate only | | | | 5 | 5 | 5 |
| Q8GUK8_ARATH | At5g10060 | At5g10060 | 13,86 | 51955 | Eluate only | | | | 5 | 5 | 5 |
| TBA2_ARATH | Tubulin alpha-2 chain | TUBA2 | 14,67 | 49541 | Eluate only | | | | 4 | 5 | 5 |
| A0A1P8BFF0_ARATH | Acyl-CoA binding protein 5 | ACBP5 | 34,58 | 73292 | Eluate only | 7 | 9 | 0 | 22 | 38 | 17 |
| HS901_ARATH | Heat shock protein 90-1 | HSP90-1 | 17,86 | 80635 | Eluate only | 2 | 3 | 0 | 13 | 14 | 10 |
| AROD3_ARATH | Arogenate dehydratase 3, chloroplastic | ADT3 | 26,42 | 46102 | Eluate only | 1 | 1 | 0 | 8 | 9 | 8 |
| PATL1_ARATH | Patellin-1 | PATL1 | 18,32 | 64046 | Eluate only | 2 | 2 | 0 | 9 | 12 | 8 |
| A1L4Y1_ARATH | At3g10350 | At3g10350 | 37,71 | 44754 | 38,00 | 1 | 1 | 1 | 18 | 38 | 38 |
| NIR_ARATH | Ferredoxin--nitrite reductase, chloroplastic | NIR1 | 42,32 | 65505 | 34,00 | 1 | 1 | 1 | 26 | 34 | 34 |
| DCE2_ARATH | Glutamate decarboxylase 2 | GAD2 | 42,11 | 56141 | 20,50 | 4 | 5 | 2 | 36 | 63 | 41 |
| SECA1_ARATH | Protein translocase subunit SECA1, chloroplastic | SECA1 | 16,24 | 115183 | 20,00 | 1 | 1 | 1 | 19 | 20 | 20 |
| PR40C_ARATH | Pre-mRNA-processing protein 40C | MED35C | 16,17 | 92807 | 17,00 | 1 | 1 | 1 | 14 | 17 | 17 |
| A0A178UKA9_ARATH | Uncharacterized protein | AXX17_At5g10570 | 36,87 | 48691 | 15,00 | 1 | 1 | 1 | 14 | 15 | 15 |
| F4K0R0_ARATH | Transducin family protein / WD-40 repeat family protein | MOP10.11 | 17,11 | 120063 | 14,00 | 1 | 1 | 1 | 14 | 14 | 14 |
| CRWN1_ARATH | Protein CROWDED NUCLEI 1 | CRWN1 | 31,1 | 129093 | 13,25 | 4 | 4 | 4 | 44 | 53 | 53 |
| A0A178UHM4_ARATH | Uncharacterized protein | AXX17_At5g00220 | 24,63 | 52331 | 13,00 | 1 | 1 | 1 | 12 | 13 | 13 |
| A0A178URW9_ARATH | Uncharacterized protein | AXX17_At5g30360 | 33,9 | 26061 | 12,00 | 1 | 1 | 1 | 8 | 12 | 12 |
| A0A178U844_ARATH | Signal recognition particle subunit SRP68 | AXX17_At5g61470 | 31,24 | 68833 | 11,50 | 2 | 2 | 2 | 20 | 23 | 23 |
| ALFC5_ARATH | Fructose-bisphosphate aldolase 5, cytosolic | FBA5 | 25,7 | 38294 | 11,00 | 2 | 2 | 1 | 9 | 12 | 11 |
| CAP2_ARATH | Putative clathrin assembly protein At2g25430 | At2g25430 | 13,02 | 72084 | 10,00 | 1 | 1 | 1 | 9 | 10 | 10 |
| DCE4_ARATH | Glutamate decarboxylase 4 | GAD4 | 38,74 | 56005 | 10,00 | 6 | 7 | 1 | 27 | 49 | 10 |
| Q9SK61_ARATH | Expressed protein | At2g20410 | 32,74 | 37569 | 10,00 | 1 | 1 | 1 | 9 | 10 | 10 |

**Table S2**: **NADKc-1 kinetic parameters**

| Substrate varied | Constant substrate | $K_M$ (µM) | $k_{cat}$ ($s^{-1}$) | $k_{cat}/K_M$ ($\mu M^{-1}.s^{-1}$) |
|---|---|---|---|---|
| ATP | $NAD^{+a}$ | 203(±30) | 41(±2) | 0.2 |
| CTP | id. | 283(±70) | 42(±1) | 0.15 |
| GTP | id. | 522(±135) | 26(±1) | 0.05 |
| UTP | id. | 207(±26) | 29.5(±2) | 0.14 |
| $NAD^+$ | $ATP^b$ | 147 (±17) | 42(±2) | 0.28 |
| NADH | ATP | - | - | - |
| NAAD | ATP | - | - | |

[a]ATP 8 mM; [b]$NAD^+$ 10 mM. Values represent the average of three independent measurements. Id: as above, -: no activity detected.

**Table S3. Comparison of CaM-dependent NADK activity in different photosynthetic organisms**

| | *A. thaliana* | *M. polymorpha* | *K. flaccidum* | *C. reinhardtii* |
|---|---|---|---|---|
| *Total activity*[a] | 31.2(±2.8) | 6.9(±0.7) | 5.8(±0.3) | 24.6(±1.4) |
| *CaM/Ca$^{2+}$-independent activity*[b] | 1.0(±0.3) | 1.0(±0.1) | 1.8(±0.3) | 29.7(±2.7) |
| *CaM/ Ca$^{2+}$- dependent activity*[c] | 30.2 | 5.9 | 4.0 | - |

NADK activity was measured in soluble protein extracts from *A. thaliana*, *M. polymorpha*, *K. flaccidum* and *C. reinhardtii as detailed in methods.* Activities are expressed in nmol/h/mg protein. [a]NADK activity measured in the presence of CaM/Ca$^{2+}$ represents CaM-independent plus CaM–dependent activity; [b]NADK activity measured in the presence of trifluoroperazine (CaM inhibitor) represents CaM-independent NADK activity. [c]CaM/Ca$^{2+}$ activity is the difference between total NADK activity and CaM/Ca$^{2+}$-independent activity. NADK activity in C. *reinhardtii* is independent on CaM/Ca$^{2+}$ (the difference is within experimental error).

**Table S4: Primers used in this study**

In red: sequences for recombination reactions by BP clonase into pDONR221.

Uppercase: restriction enzyme sites.

**Primers for cloning the NADKc-1 protein sequence for enzymatic, localization and complementation studies:**

| construct | Forward primer | Reverse primer |
|---|---|---|
| pet28(b)-6-HIS-NADKc-1 | gacttttcaaaccaaCATATGgtgaaaccc ttaggagaag | caaattcaggaagaagaCTCGAGcaaaggtc tagttc |
| pet28(b)-6-HIS-Δ38NADKc-1 | gctgtagcagctCATATGgccggagaatta ctc | caaattcaggaagaagaCTCGAGcaaaggtc tagttc |
| 35S::NADKc-1*N*R::YFP | GGGGACAAGTTTGTACAAAAAAGCAGGCTT gatggtgaaacccttaggagaag | GGGGACCACTTTGTACAAGAAAGCTGGGTgg acatctttataccccatg |
| 35S::YFP::NADKc-1 and 35S::NADKc-1 (complementation) | GGGGACAAGTTTGTACAAAAAAGCAGGCTT gatggtgaaacccttaggagaag | GGGGACCACTTTGTACAAGAAAGCTGGGTtc aattttttggttgggccttttcgattctcc |

**Primers for genotyping:**

| line | Forward primer | Reverse primer | LB primer |
|---|---|---|---|
| SALK_130871 | tgttaagcaaatatgtgggcc | acaaatgatcgaaatggcaag | LBb1.3 attttgccgattttcggaac |
| GABI_311H11 | taaggaagaaggacgggacttc | tcaagatcgaccaaagcattag | LBgabi-kat atattgaccatcatactcattgc |

**Primers for gene expression studies (qPCR):**

| | Forward primer | Reverse primer |
|---|---|---|
| At1g04280 | aacgggtcagcagtctcaac | gggcctcgatggaactaaca |

### Supplemental Material and Methods

#### Partial purification of native NADKc-1

To purify native NADKc-1, Col-0 Arabidopsis plants seeds were sterilized, sown in Murashige and Skoog liquid media (2.2 g $L^{-1}$) supplemented with 0.5 g $L^{-1}$ sucrose and grown under continuous light (60 µE) in 250-mL flasks and under agitation (125 rpm). 7-day-old plants were rinsed with water, frozen in liquid nitrogen and ground with a mortar and pestle. The powder was suspended in Buffer A (50 mM Tris-HCl pH 7.5, 400 mM KCl, 3 mM $MgCl_2$, 1 mM EGTA, 0.5 mM EDTA, 1 mM DTT, 1 mM $NAD^+$, 10 µM leupeptine, 10 µM E64, 1 mM PMSF, 1 mM benzamidine and 5 mM ε-aminocaproic acid). The crude extract was centrifuged (15,000 g, 30 min, 4 °C) and the recovered soluble protein extract (432 mg, 72 mL, 0.74 µmol $h^{-1}$ $mg^{-1}$) was precipitated with 50% ammonium sulfate (1 h, 4 °C) and centrifuged (16,000 rpm, 20 min). The protein pellet was then suspended in 32 mL Buffer A (30 mL, 270 mg, 1.1 µmol $h^{-1}$ $mg^{-1}$) and loaded onto a DEAE column (60 mL resin) equilibrated with Buffer A. AtCaM1/$Ca^{2+}$-dependent NADK activity was detected in the flow-through (120 mL, 84 mg, 1.8 µmol $h^{-1}$ $mg^{-1}$). NaCl (3.5 M) was added to this fraction, and the protein solution was loaded onto a Butyl-sepharose column (12 mL resin) equilibrated with Buffer B - NaCl (50 mM Tris-HCl pH 7, 3.5 M NaCl, 100 mM KCl, 3 mM $MgCl_2$, 1 mM DTT). Elution was performed by applying a linear gradient (60 mL) from 0 to 100% Buffer B-ethane diol (50 mM Tris-HCl pH 7.5, 30% ethane diol (v/v), 100 mM KCl, 3 mM $MgCl_2$, 1 mM DTT). Eluted fractions containing AtCaM1/$Ca^{2+}$-dependent NADK activity were supplemented with 1 mM $NAD^+$ and immediately frozen. Fractions were pooled (5.6 mg, 2.32 µmol $h^{-1}$ $mg^{-1}$, 8 mL) before loading onto a 1-mL CaM-sepharose column (Stratagen) equilibrated with buffer C (50 mM Tris pH 7.5, 10% (v/v) Glycerol, 1 mM DTT, 2 mM $CaCl_2$). The column was washed with 10 volumes Buffer C supplemented with 500 mM NaCl. Bound proteins were then eluted with 50 mM Tris pH 7.5, 10% (v/v) Glycerol, 1 mM DTT and 5 mM EGTA. The fraction with highest activity (3 µg, 100 µl, 577 µmol $h^{-1}$ $mg^{-1}$) was used for LC-MS/MS analyses (Fig. S1).

#### Mass spectrometry-based proteomics analyses

Proteins from the eluate and flow-through fractions were stacked by performing a very short run on an SDS-PAGE gel (NuPAGE 4-12%, ThermoFisher Scientific). Proteins were revealed by staining with Coomassie blue (R250, Bio-Rad), and submitted to in-gel digestion using modified trypsin (Promega, sequencing grade), as previously described (Casabona et al., 2013).

Resulting peptides were analyzed by online nanoLC-MS/MS (UltiMate 3000 and LTQ23 Orbitrap Velos Pro, Thermo Scientific). Briefly, peptides were sampled on a 300 µm x 5 mm PepMap C18 precolumn and separated on a 75 µm x 250 mm C18 column (PepMap, Thermo Scientific). MS and MS/MS data were acquired using Xcalibur (Thermo Scientific). Peptides and proteins were identified using Mascot (version 2.6), performing concomitant searches against Uniprot (*A. thaliana* taxonomy), the classical contaminants database (in-house) and the corresponding reversed databases. Proline software (http://proline.profiproteomics.fr) was used to filter the results (conservation of rank 1 peptides, peptide identification FDR < 1% as calculated on peptide scores using the reverse database strategy, minimum peptide score of 25, and minimum of one specific peptide identified per protein group). Results from individual samples were compiled, and grouped before comparing protein groups from different samples. Proteins were considered to be enriched in the eluate if they were identified only in this sample with a minimum of five specific spectral counts, or if they were enriched at least 10-fold in this sample compared to flow-through sample. Relative quantification was performed on the basis of specific spectral counts.

**Recombinant protein expression and purification**

Full-length At1g04280 cDNA was amplified by PCR from a pYES cDNA library (Elledge et al., 1991) using the primers listed in Supplemental Table S1. The PCR products digested by Nde I and Xho I were ligated into a pet28b(+) vector digested by the same enzymes to produce the constructs named pet28(b)-6-HIS-NADKc-1 and pet28(b)-6-HIS-Δ38NADKc-1. The proteins were produced from these constructs with a 22-amino-acid N-terminal His-tag.

Rosetta2-competent cells were transformed by one or the other of these two constructs and grown at 37 °C in the presence of Chloramphenicol (34 µg.mL-1) and Kanamycin (50 $\mu g.mL^{-1}$) until Abs at 600 nm reached 0.6. IPTG (0.4 mM) was added, and growth was continued at 20 °C for 15 h. Bacteria were pelleted, suspended in Buffer D (50 mM Hepes pH 7.8, 500 mM KCl, 10% (v/v) glycerol, 50 mM L-arginine, 50 mM L-Glutamate, 1 mM $NAD^+$) supplemented with 1 mM benzamidine and 5 mM ε-aminocaproic acid, 1 mM β-mercaptoethanol, and lysed by sonication. Insoluble material was removed by centrifugation (15,000 g, 20 min, 4 °C), and soluble proteins were recovered. A much higher production level and higher activity were obtained with the Δ38-NADKc construct, therefore further work was performed with this protein. The protein extract (30 mL, 11.3 mg $mL^{-1}$) was loaded onto a Ni-NTA sepharose column (3 mL resin). After washing with buffer D supplemented with 50 mM imidazole (15 mL), the recombinant protein was eluted in buffer A supplemented with 250 mM imidazole. The imidazole concentration was reduced to 5 mM by protein-concentration cycles followed by dilution in buffer A. Pooled fractions containing NADKc-1 activity were concentrated on 10K Vivascience concentrators, aliquoted, frozen in liquid $N_2$ and stored at -80 °C. After this first purification step, the protein fraction still contained small amounts of contaminating proteins. To improve purification, Ni-NTA-sepharose fractions (2 mg, 2.2 mL) were denatured in 8 M urea and purified on Ni-NTA sepharose under denaturing conditions (6 M urea). After washing with 15 mL buffer E (25 mM Tris-Cl pH 8.0, 500 mM NaCl, 5 mM β-mercaptoethanol, 6 M urea) supplemented with 50 mM imidazole, the pure denatured protein was eluted in buffer E supplemented with 250 mM imidazole. The denatured protein (0.3 mL, 0.3 mg, lane 4, Fig. S4) was then refolded by drop-by-drop dilution in 15 mL buffer D supplemented with 1 mM DTT. The refolding reaction was performed with mixing and at room temperature. The highly pure protein (lane 5, Fig. S4) was then concentrated as described above, aliquoted, frozen in liquid nitrogen and stored at -80 °C.

*A. thaliana* Calmodulin 1 (AtCaM1, At5g37780) was purified as previously described (Dell'Aglio et al., 2013).

Proteins were quantified by the Bradford method, using bovine γ-globulin as a standard or, for purified proteins, by measuring A205nm (Scopes, 1974). CaM-binding peptide (QKVPKLKDFVMAATRKQRFERVTKDLKVKR) was synthesized by Smart Bioscience (France). The control peptide sequence was VKDTELAKKVWDFSTKLTDS (Dell'Aglio, 2013).

**Activity measurements**

NADK activity in extracts prepared from Col-0 *A. thaliana* plants, *K. flaccidum*, *M. polymorpha* and *C. reinhardtii* was measured at 25 °C as previously described (Turner et al., 2004) in the presence of a CaM inhibitor (trifluoroperazine, 50 µM) or in the presence of AtCaM1 (1 µM) and $Ca^{2+}$ (500 µM) . Recombinant NADKc activity was measured in thermostatic cuvettes (25 °C) with 20 nM enzyme in the presence of 50 mM Hepes-KOH pH 7.8, 5 mM glucose-6-phosphate, 10 mM $MgCl_2$ and variable concentrations of $NAD^+$ and ATP (or other nucleotide), 30 mU of glucose-6-phosphate dehydrogenase from baker's yeast and, where indicated in the figure legends, 50 µM $Ca^{2+}$ and/or AtCaM1. NADPH production was detected by monitoring absorbance at 340 nm ($\varepsilon_{340\ nm}$ = 6250 $M^{-1}\ s^{-1}$). Kinetic data reported in Table 1 were calculated by fitting enzymatic data to the appropriate theoretical equations using the Kaleidagraph program (Synergy Software, Reading, PA, USA). Tight-binding of AtCaM1 onto NADKc-1 was fitted using Eqn. 1:

$$v/[E]0 = k\text{cat}\frac{([\text{CaM}] + k\text{d} + [\text{E}]0) - \sqrt{([\text{CaM}] + k\text{d} + [\text{E}]0)^2 - 4 * k\text{d} * [\text{E}]0}}{2[\text{E}]0}$$

(Eqn.1)

where *k*cat is the catalytic constant for NADKc, [CaM] is the total CaM concentration, *k*d is the dissociation constant for the interaction between CaM and NADKc-1, and [E]0 is the total NADKc-1 concentration.

**Subcellular localization by confocal microscopy**

The NR of the protein (amino acids: 1-60) or the entire protein sequence was amplified by PCR from pet28(b)-6-HIS-NADKc-1 using the primers listed in Supplemental Table S1. The amplified PCR products were cloned into the Gateway-adapted vector pDONR211 by BP recombination, to produce pDONR:NADKc-1*NR* and pDONR:NADKc-1STOP constructs.

These vectors were then recombined with pB7YWG2,0 and pB7WGY2,0 vectors (Karimi et al., 2007), respectively, to generate pB7YWG2:NADKc-1*NR*:YFP and pB7WGY2:YFP:NADKc-1 through the LR recombination reaction. Both vectors were transformed into *Agrobacterium tumefaciens* C58 1 and used for tobacco leaf infiltration.

Transient expression in *Nicotiana benthamiana* var. Xanthi was performed by infiltrating leaves with a suspension of *A. tumefaciens* harboring pB7YWG2:NADKc-1*NR*:YFP and a mitochondrial marker (pSu9–RFP) (Michaud et al., 2014) or pB7WGY2:YFP:NADKc-1, and the silencing suppressor P19 protein.

Transformed samples were observed by confocal microscopy on a LSM 800 confocal microscope equipped with a Plan-Apo 63X/1.4 oil DIC lens, two GaAsP detectors and one Airyscan detector. YFP and RFP fluorophores were excited at 488 and 558 nm, respectively, and emission signals were measured at 488–560 nm for YFP, and at 568-611 nm for RFP.

**Measurement of photosynthetic parameters**

Maximum quantum yield of photosystem II ($F_V/F_M$), relative electron transport rate (ETR) and non-photochemical quenching (NPQ) were determined using a chlorophyll fluorescence imaging system (SpeedzenIII, JbeamBio, France) Fv/Fm: (Fm-$F_O$)/Fm; ETR = (($F_M$' – $F_S$)/$F_M$'); NPQ = ($F_M$ – $F_M$')/$F_M$'; where $F_M$ = maximum fluorescence; $F_O$ = minimum fluorescence; $F_V$ =variable fluorescence ($F_V = F_M - F_O$) in dark-adapted state (Maxwell and Johnson, 2000). $F_M$', maximum fluorescence in the light; $F_S$, steady-state chlorophyll fluorescence. Plants were dark adapted for at least 30 minutes prior to measurement.

**Mutant complementation**

To complement the SALK_006202 NADKc-1 mutant line, the entire NADKc-1 protein sequence was amplified by PCR from pet28(b)-6-HIS-NADKc-1 using the primers listed in Supplemental Table S1. The amplified PCR product was cloned into the Gateway-adapted vector pDONR211 by BP recombination, thus generating pDONR:NADKc-1STOP. This vector was then recombined with pB2GW7,0 (Karimi et al., 2007) to generate pB2GW7:NADKc-1 by the LR recombination reaction. This vector was transformed into *A. tumefaciens* C58 and used for floral dipping transformation. Complemented lines were screened based on their BASTA-resistance and AtCaM1/$Ca^{2+}$- dependent NAD kinase activity.

Lines for which levels of AtCaM1/$Ca^{2+}$-dependent NAD kinase activity were similar to wild-type plants were further analyzed.

**Oxidative burst measurements**

*A. thaliana* plantlets were grown for seven days in continuous light (Benamar A, 2013) and immunity-related accumulation of $H_2O_2$ following stimulation with flg22 (1 µM) was analyzed as previously described (Bisceglia & Savatin, 2015). Luminescence measurements were performed with a microplate reader SPARK 10M (TECAN) using 96-well microtiter plates (flat bottom, white, Greiner). For each plant type, results are expressed as the average luminescence obtained for 4 wells (*i.e.*, 120 plantlets).

**Molecular Phylogenetic analysis by the Maximum Likelihood method**

Protein sequences of NADKc-like proteins were retrieved from the Phytozome website (Goodstein et al., 2012), with some exceptions. The sequences of *U. mutabilis* and *C. braunii* were retrieved from the ORCAE project website (Sterck et al., 2012) and those of *S. minuta*, *E. fimbriata*, Mougeotia sp. and Spirogyra sp. were retrieved from the One KP project website (Matasci et al., 2014). The evolutionary history of NADKc-1 was inferred by using the Maximum Likelihood method, based on the JTT matrix-based model (Jones et al., 1992). The tree with the highest log-likelihood (-31386.44) is shown. The percentage of trees produced in which the associated taxa clustered together is shown next to the branches. Initial tree(s) for the heuristic search were obtained automatically by applying Neighbor-Join and BioNJ algorithms to a matrix of pairwise distances estimated using a JTT model, and then selecting the topology with the highest log-likelihood value. A discrete Gamma distribution was used to model evolutionary rate differences between sites (5 categories (+G, parameter = 0.9891)). The tree is drawn to scale, with branch lengths based on the number of substitutions per site. The analysis involved 45 amino acid sequences. There were a total of 1196 positions in the final dataset. Evolutionary analyses were 1 performed in MEGA7 (Kumar et al., 2016).
